# Tuned Fitness Landscapes for Benchmarking Model-Guided Protein Design

**DOI:** 10.1101/2022.10.28.514293

**Authors:** Neil Thomas, Atish Agarwala, David Belanger, Yun S. Song, Lucy J. Colwell

## Abstract

Advancements in DNA synthesis and sequencing technologies have enabled a novel paradigm of protein design where machine learning (ML) models trained on experimental data are used to guide exploration of a protein fitness landscape. ML-guided directed evolution (MLDE) builds on the success of traditional directed evolution and unlocks strategies which make more efficient use of experimental data. Building an MLDE pipeline involves many design choices across the design-build-test-learn loop ranging from data collection strategies to modeling, each of which has a large impact on the success of designed sequences. The cost of collecting experimental data makes benchmarking every component of these pipelines on real data prohibitively difficult, necessitating the development of *synthetic* landscapes where MLDE strategies can be tested. In this work, we develop a framework called SLIP (“Synthetic Landscape Inference for Proteins”) for constructing biologically-motivated synthetic landscapes with tunable difficulty based on Potts models. This framework can be extended to any protein family for which there is a sequence alignment. We show that without tuning, Potts models are easy to optimize. In contrast, our tuning framework provides landscapes sufficiently challenging to benchmark MLDE pipelines. SLIP is open-source and is available at https://github.com/google-research/slip.

## 1 Introduction

Directed evolution (DE) [1–3] has revolutionized bioengineering, enabling the development of proteins with novel function across industries including food, chemicals, and therapeutics [2, 4, 5]. Part of the power of directed evolution is its simplicity: synthesize a set of variants, screen them for the desired function, induce mutagenesis in the top variants that passed the screen and repeat. This selective pressure pushes the variants towards the desired activity without requiring any prior knowledge of how mutations affect function. We can view DE as a genetic algorithm exploring a high-dimensional *landscape*, where higher points correspond to greater levels of desired activities. DE works under the assumption that functional landscapes are relatively smooth [3, 6], that is, combining high-performing mutations results in further improved variants.

With sufficient experimental throughput, DE can successfully climb smooth fitness landscapes. However, testing a set of variants for activity is a costly and time-consuming process, requiring specialized laboratory expertise. For example, experimental throughput to test the turnover rate of an enzyme can be on the order of tens of variants per round over a 10-round campaign [7]. Each inactive design comes at a great cost, not only because of the cost of the single negative result, but also comes at an opportunity cost as the inactive variant cannot be leveraged for designs in future rounds of DE. Thus, a practitioner wants to make efficient use of their experimental budget by carefully curating the designed variants. Guiding the designs can be done with *rational design*, which uses an understanding of the protein’s structural and functional properties to avoid poor designs. However, these properties are often unknown: an arbitrary wildtype starting sequence may not have any associated structure [8], or any characterization besides its homology to other sequences. To gain the understanding that would enable rational design would require experimental characterization which could be even more costly than the rest of the protein design campaign.

Effectively exploring rough (i.e., not smooth) landscapes with limited experimental throughput is a challenge for DE [9–11]. Thankfully, the confluence of advancements in DNA synthesis (bespoke oligonucleotide sequences), DNA sequencing (high throughput screens), and machine learning have enabled a new paradigm of machine learning guided directed evolution (MLDE) [12–15] which can address the challenge of exploring rough landscapes with limited data. In each round of MLDE, a set of (variant, activity) pairs are collected, which a practitioner can use to train a genotype-phenotype model to predict the effect of variants. In the next round, that model can be used to propose candidates (see Figure 1 in [12]). MLDE has been successfully applied to designing proteins, such as AAV capsids with preferential delivery to specific organs in the body [16, 17], or increasing the fluorescence of GFP [18].

**Fig. 1:**
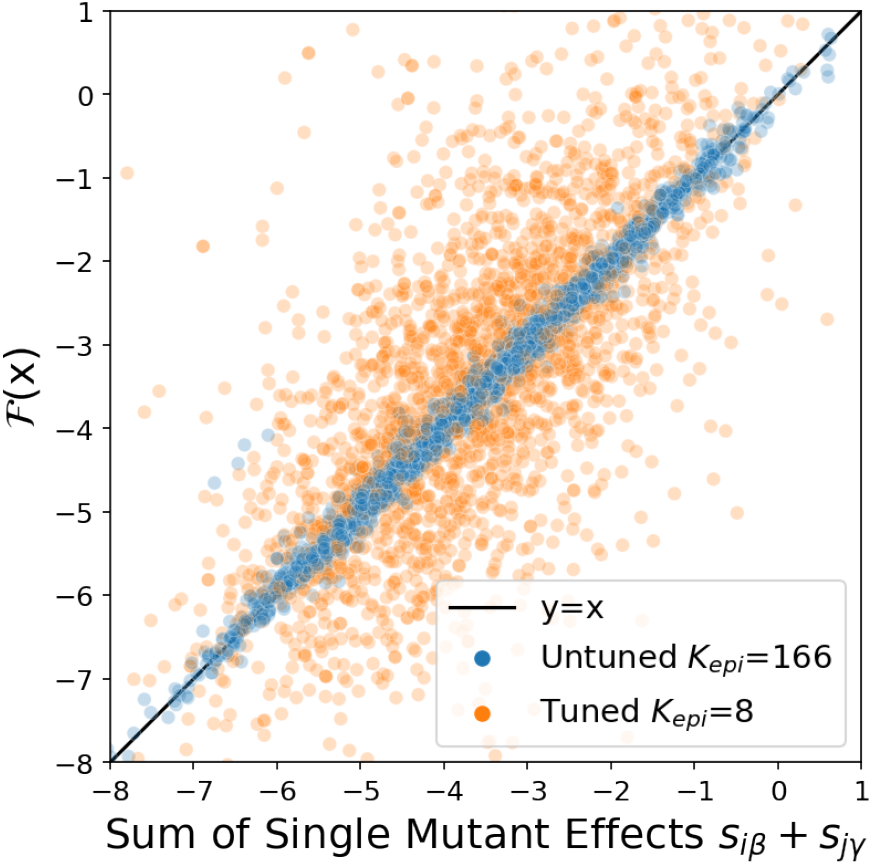
Tuning the epistatic horizon increases the ruggedness of the resulting landscape. Fitness ℱ(**x**) of 5000 variants with 2 mutations (centered so that ℱ(**x**_0_) = 0). The untuned fitness landscape with epistatic horizon *K*_epi_ = 166 (blue) is roughly linear, while the tuned fitness landscape with epistatic horizon *K*_epi_ = 8 (orange) exhibits more ruggedness. This landscape is derived from the alignment for PDB id 3er7. Note that the untuned epistatic horizon is greater than the length of the protein, *K*_epi_ = 166 > 118 = *L*.

Despite early successes, there is no consensus on best practices for MLDE. There are many decisions involved in the design of an MLDE pipeline. What data should be collected? Which model class should be used? What optimization objective should be used for model training? How should models be selected? How should proposals be generated using a trained model? How does one trade off between “exploiting” model confidence and “exploring” regions of model uncertainty when synthesizing the next round of designs [19,20]? Each choice interacts nonlinearly with every other choice, complicating the meta-optimization problem.

Running experiments is expensive, so it is essential to test the pipeline before using it in a new design campaign. One approach is to use *empirical landscapes* from publicly available experimental datasets. However, these datasets are limited by the prohibitive cost of producing enough high-quality data to benchmark an MLDE pipeline across multiple rounds. Datasets like [13, 17, 21–23] curate the highest quality protein function datasets from Deep Mutational Scanning [24], but focus on one- or two-mutation regions around a wildtype sequence [25]. Other landscapes contain all possible multi-mutants, but only cover a small portion of the overall protein (4 positions of the binding domain (B1) of protein G [26]), or test a limited allele vocabulary [27].

In parallel with deeply characterized empirical landscapes, *synthetic landscapes* are being explored as testing grounds for MLDE [20, 28]. Synthetic landscapes are defined by a software function that can be queried for any sequence of interest [10, 19, 28–32]. In this paper, we propose a specific synthetic landscape with two key properties:

− Properties **grounded** in the statistics of real protein families.
− **Tunable, interpretable** difficulty to match a range of plausible optimization landscapes.

To obtain these properties, we introduce and validate an MLDE benchmarking framework called SLIP: “Synthetic Landscape Inference for Proteins.” SLIP is a set of synthetic fitness landscapes based on Potts models [33–37], combined with utilities for tuning the landscape difficulty. SLIP is open-source and is available at https://github.com/google-research/slip.

## 2 Background and Related Work

We define a *fitness landscape* ℱ as a scalar-valued function over sequences **x** of length *L* with *A* alleles at each position. In practice, “fitness” is used to refer to a molecular phenotype (e.g., fluorescence) or organismal fitness (e.g., reproductive rate). Empirical landscapes measure the underlying fitness through a noisy observation process. Our synthetic fitness landscapes refer to the underlying, noiseless quantity.

We will consider a sequence design process which starts at “wildtype” **x**_0_, which has allele *a*_*i*_ at site *i*. The goal of the design process is to *maximize* the fitness (as opposed to the minimization of the loss function that occurs in traditional supervised learning). Therefore, we focus on the fitness *gain* ℱ(**x**) − ℱ(**x**_0_). A successfully designed sequence will, at a minimum, display ℱ(**x**) − ℱ(**x**_0_) > 0.

### 2.1 Epistasis and Landscape Ruggedness

In order to test an MLDE pipeline, we need landscapes that cannot be effectively navigated without guidance from an ML model. Thus it is useful to tune the nonlinearity (or “difficulty”) of a landscape. Two basic quantities we can use to understand the difficulty of task are 1) the *single-mutant effects* and 2) *pairwise epistasis*.

#### Single-mutant effects

Letting **x**_*iβ*_ denote the sequence obtained by mutating the wildtype allele at site *i* to allele *β*, the single mutant effect *s*_*iβ*_ is defined as:

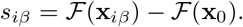

Landscapes with many *adaptive* single mutations (site-allele pairs where *s*_*iβ*_ > 0) tend to be easier to optimize.

#### Pairwise epistasis

If the fitness of multi-mutants was given by the sum of single-mutant effects, then the landscape would be linear and easy to optimize – simply combining adaptive single mutants would lead to good sequences **x** with high values of ℱ(**x**). Nonlinear interactions between mutations make optimization more difficult. These nonlinear effects are known as *epistasis*, a pervasive property in empirical landscapes [9, 38–42]. For two mutations *a*_*i*_ →*β* at site *i* and *a*_*j*_ → *γ* at site *j*, we can define the *pairwise epistasis ϵ*_*iβ,jγ*_ by

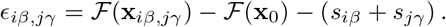

In other words, pairwise epistasis is the part of the fitness difference between **x**_*iβ,jγ*_ and **x**_0_ which cannot be explained by the single mutants. Deleterious epistatic interactions between adaptive single mutants (known as *reciprocal sign epistasis*) make it more difficult to combine good individual mutations to obtain good multi-mutants, confounding a linear model of the landscape.

Note that both pairwise epistasis and the single-mutant fitness differences must be defined relative to a reference sequence - in our case, the wildtype **x**_0_.

### 2.2 Synthetic Landscapes

Synthetic landscapes are designed so that ℱ can be evaluated quickly for arbitrary sequences, even if computing all *L*^*A*^ values is prohibitive (in memory or time). Many of these landscapes were originally designed to study evolutionary processes [10, 32, 43].

More recently, synthetic landscapes have been applied to benchmark sequence design algorithms [20, 28, 31]. We can divide synthetic landscapes into four broad, overlapping classes: supervised neural models, biophysical models, random models, and graphical models, briefly described below.

#### Supervised neural models

are trained on experimental data to predict a regression output [44–47]. Of particular note is the structural score given by AlphaFold2, which can be used as an optimization objective to find sequences likely to fold into a desired structure [46,48]. If the model is sufficiently good, model outputs can be used as a synthetic replacement for the experimental target. Neural models are fast to evaluate with a single forward pass. However, they can exhibit pathological behavior when used as optimization objectives, giving high scores to unrealistic sequence [49, 50] or giving outsize influence to irrelevant parts of the sequence [51]. While trained neural models can exhibit high levels of ruggedness [52], it is not straightforward to tune the optimization difficulty of a neural landscape.

#### Biophysical models

explicitly model the energetic interactions in the protein. Prominent examples of biophysical models are ViennaRNA [44] and Rosetta [53], which provide a score for an input sequence representing the free energy of the folded structure at equilibrium. These models return globally characterized landscapes without unexpected pathologies. However, they require a computationally expensive optimization procedure to report the free energy minimum for each query sequence. Since physical assumptions like physical constants and potential energy functions are baked into the model, there is no principled tuning procedure to make the landscapes more difficult to optimize.

#### Random models

generate a landscape by drawing a random function on the configuration space {1, …, *A*}^*L*^ with a particular distribution. The NK-model explicitly models order-*K* interactions for a sequence of length *N* to provide tunably rugged fitness landscapes which exhibit high-order correlations [29, 30]. The generalized NK-model, which explicitly models sparse blocks of interacting positions, has been shown to reflect the sparsity of empirical fitness function when conditioned on real structures [54]. Distance-dependent models [11] have interactions at all orders, and are defined by fixing a functional form for the covariance between sequences as a function of genetic distance. All of these models have been applied to study evolutionary dynamics, and are typically not fit to data.

#### Graphical models

explicitly represent the interactions between positions in the sequence as edges in a graph. Profile HMMs do not model epistasis, and only model first-order interactions (i.e., amino acid distributions at aligned positions). Despite this, they serve as powerful protein family classifiers [55], and HMM likelihoods have been used as synthetic optimization objectives [28]. Profile HMMs can also flexibly handle insertions and deletions, and do not require aligned sequences as input. We describe in detail below a graphical model known as a Potts model.

### 2.3 Potts Models of Protein Families

#### Definition

For a family of proteins of length *L* with *A* possible alleles at each position, a Potts model defines a probability distribution over sequences in the family as

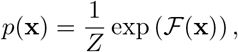

where ℱ is a negated statistical energy, and the partition function *Z* is a normalization constant such that the *p*(**x**) sum to 1. In a Potts model, for an input one-hot encoded sequence **x** ∈ ℝ^*L*×*A*^, ℱ is given by the sum over the marginal effects and pairwise interactions:

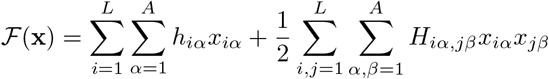

where **h** = (*h*_*iα*_) is a tensor of dimension *L* ×*A* representing marginal terms and **H** = (*H*_*iα,jβ*_) is a symmetric tensor of dimension *L* × *L* × *A* × *A* representing pairwise coupling terms. The parameters of a Potts model can be fit using a set of aligned sequences; see Supplementary Material A.2 for details.

#### Modeling coevolution

There has been extensive work establishing that Potts models learn statistics grounded in protein structure and function. Potts models are useful as unsupervised structure predictors [35–37, 56], and are competitive with neural unsupervised structure predictors [57] on families with large, diverse alignments. The statistical energy of protein variants scored by a Potts model has been shown to correlate well with empirical fitness [58]. When the statistical energy is included as an additional feature to a regression model, it has been shown to improve predictive performance on empirical landscapes [25]. Used as generative models, Potts models have been shown to propose functional variants of a given protein target [59]. Synthetic sequences evolved *in silico* on a Potts landscape have been shown to correlate with summary statistics with *in vitro* evolved sequences [60]. The parameters of the Potts model can also be used as input featurizations that improve performance on downstream tasks [61, 62].

## 3 Methods: Tuned Quadratic Landscapes

We can use the statistical energy of the Potts model as the fitness function ℱ to define a synthetic landscape. In [28], for example, the authors introduced a “PDB-Ising” synthetic fitness landscape, which combined the contact map of a protein with standard pair potentials for amino acid substitution to create a simple *quadratic landscape* with pairwise interactions. We will instead derive model parameters from alignment data, which has the advantage that the resulting synthetic landscape exhibits correlations grounded in the coevolutionary couplings in that family.

While useful in their own right, Potts models derived from alignment data are not challenging synthetic landscapes, as they can be optimized by combining top mutations one at a time (akin to an *in silico* DE algorithm). We quantify these shortcomings in Section 5. To benchmark the performance of MLDE pipelines on more difficult optimization scenarios, we require synthetic landscapes where strategies guided by nonlinear models can substantially outperform those guided by linear models. In what follows, we develop a framework for tuning quadratic landscapes to a desired level of difficulty.

### 3.1 Tuning Model Statistics

For a fitness function defined by a Potts model, the single-mutant and pairwise epistasis terms can be written explicitly in terms of the model parameters **h** and **H**:

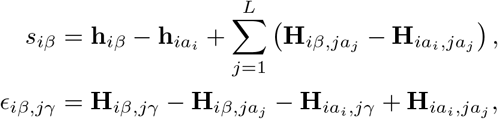

where again *a*_*i*_ is the allele of the wildtype **x**_0_ at site *i*. Note that single-mutant effect *s*_*iβ*_ depends on both the linear and quadratic parameters of the Potts model. See Supplementary Material B for a detailed derivation.

Once we reparameterize the Potts model in terms of the single-mutant and pairwise epistasis terms, the fitness decomposes into the form

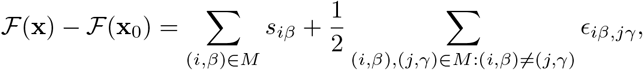

where *M* is the set of mutations in **x** encoded as site-allele pairs (*i, β*).

Introducing shift (*µ*_*s*_, *µ*_*ϵ*_) and scale (*λ*_*s*_, *λ*_*ϵ*_) parameters for single-mutant and epistatic terms, we can parameterize a family of tuned fitness functions 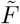 with parameters 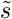 and 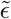 given by

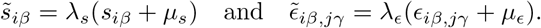

The four parameters (*µ*_*s*_, *µ*_*ϵ*_, *λ*_*s*_, *λ*_*ϵ*_) allow for the mean and variance of both 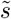 and 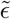, taken over position-allele pairs, to be independently tuned. This allows us the flexibility of changing the difficulty of the landscape (e.g., by making epistasis more negative on average) while maintaining much of the structure of the original problem (e.g., preserving the coevolutionary couplings).

### 3.2 Epistatic Horizon

For untuned landscapes, the single-mutant fitness effects approximate the double-mutant fitness effects well (Figure 1, blue), meaning the landscape is very linear. If combining random adaptive single mutants (mutations where *s*_*iβ*_ > 0) is not a viable strategy, a naïve design strategy will struggle. For example, in Figure 1, points with *s*_*iβ*_ + *s*_*jγ*_ > 0 but ℱ(**x**) − ℱ(**x**_0_) *<* 0 (bottom right, in orange) would be candidate proposals by a naïve algorithm that would fail an experimental screen on a rugged landscape. One way to quantify the linearity (or non-linearity) of the landscape is to define an “epistatic horizon” *K*_epi_ – the number of mutations after which the linear approximation breaks down. With an eye towards tuning optimization difficulty, we define *K*_epi_ as follows.

Let 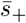 be the average of *s*_*iβ*_ over adaptive singles, and 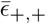 the mean epistatic effect over random adaptive pairs. Then, the sequence design problem is difficult if the average interaction between individually good mutations is negative: 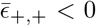. Taking *K* random adaptive mutations, the average change in fitness for a *K*-mutant **x**_*K*_ is

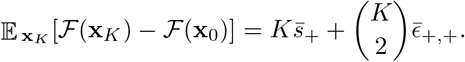

As *K* increases, the relative effect of epistasis grows relative to the single-mutant effects. We can compute a crossover value when 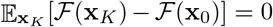. Motivated by this example, we define the *epistatic horizon K*_epi_ as the non-zero solution to the equation:

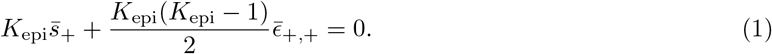

This definition suggests two ways in which synthetic landscapes can fail to be difficult for MLDE:

− *K*_epi_ *> L*. In this case, the landscape is relatively linear at all relevant length scales. Combining adaptive single mutants leads to adaptive multi-mutants even for a large number of mutations.
− *K*_epi_ *<* 0. In this case, 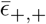 is positive; that is, on average, adaptive single-mutants combine to be *better* than the sum of their parts. In this case, combining adaptive single-mutants also leads to adaptive multi-mutants, since typical pairs will interact positively with each other.

All the untuned landscapes we studied have *K*_epi_ *> L*, and many also have *K*_epi_ *<* 0; see Figure 1 and Supplementary Table S1. This means that the untuned landscapes are unsuitable for testing MLDE pipelines as is.

However, we can use tuning parameters to adjust *K*_epi_ and generate landscapes which are more non-linear and harder to optimize (Figure 1, orange). In the next section, we derive a tuning procedure which adjusts *K*_epi_ while leaving many other statistics of the fitness landscape fixed, thereby allowing us to benchmark MLDE pipelines.

### 3.3 Tuning Procedure

With four free parameters (two shift and two scale parameters), we can introduce additional constraints on the fitness landscape. First, we normalize the landscape such that the single-mutant effects have unit variance by setting *λ*_*s*_ = *σ*(*s*)^−1^, where *σ*(*s*) is the standard deviation of the {*s*_*iβ*_}. Second, we preserve the fraction of adaptive single mutants 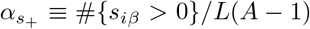 by setting *µ*_*s*_ = 0. Finally, to preserve the relative magnitudes (i.e., the ratios) of the pairwise epistatis terms, we set *µ*_*ϵ*_ = 0. This leaves *λ*_*ϵ*_ free. By fixing a target *K*_epi_, we can use Equation (1) to solve for *λ*_*ϵ*_.

To summarize: given a target epistatic horizon *K*_epi_ > 0, the tuning parameters are given by equations:

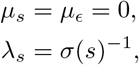

and

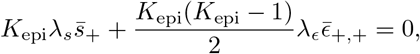

where 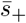 and 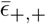 are the values for the untuned landscape. Note that other tunings are possible for a given *K*_epi_. This specific tuning scheme was chosen to preserve as much of the co-evolutionary structure of the Potts model as possible.

## 4 Methods: *In silico* Validation

To validate that our tuned synthetic landscapes are sufficiently difficult, we aim to create landscapes where nonlinear models outperform linear (naäve) models on sequence design tasks. To do so, we design an experimental framework which evaluates how effective linear and nonlinear models are at using training data to accurately rank design candidates.

We use the following procedure for each untuned landscape; see Figure 2 for a schematic:

**Fig. 2:**
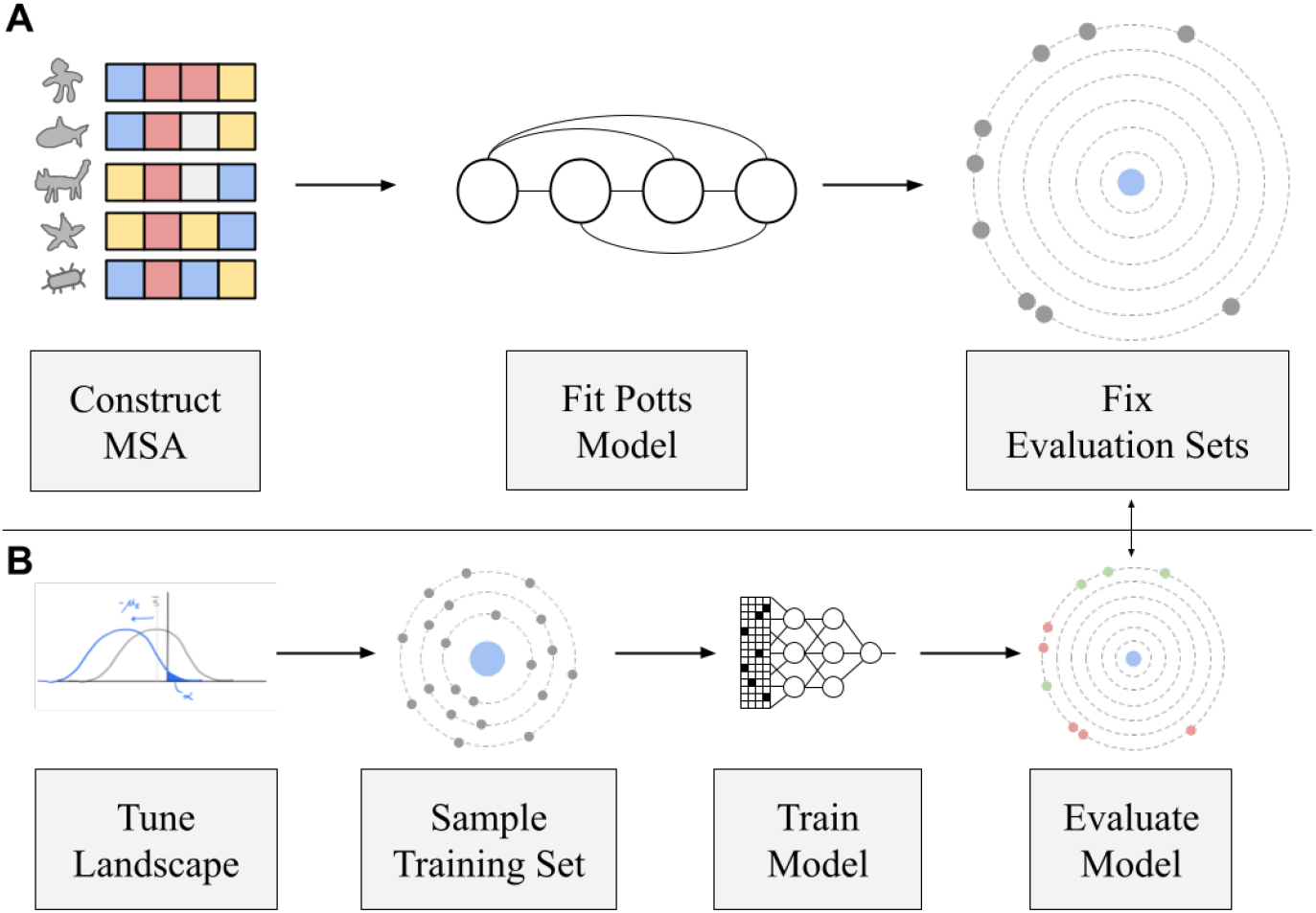
*In silico* validation workflow. Panel A shows the tasks that are performed once for each PDB ID. After training a Potts model on an aligned set of sequences, we derive a set of evaluation sequences. Panel B shows the tasks that are performed for each replicate of the regression experiment: we tune the landscape, sample a set (*x, y*) ∈ *D* of training sequences *x* with their associated synthetic fitnesses *y*, train a model on ∈ *D*, and then evaluate the model predictions on the evaluation sets. The evaluation sets are fixed for all landscapes derived from the same untuned Potts initialization.

1. Tune the epistatic horizon to *K*_epi_ = 2^𝓁^ for 𝓁 ∈ {1, 2, …, 10} as in Section 3.3. Center at ℱ(**x**_0_) = 0.
2. Sample a dataset *D* = {(**x**, *y*)} of sequences **x** with their associated fitness *y*, where |*D*| = 5000. Each sample (**x**, *y*) is obtained by sampling a number of mutations from the wildtype **x**_0_ uniformly at random from {1, 2, 3} and then sampling a variant uniformly at random at the selected distance.
3. Train Ridge and convolutional neural network (CNN) regression models across many different hyperparameter choices.
4. For the best performing model of each type, compute a paired performance metric on the evaluation set.

### 4.1 Untuned Landscapes

To select a suitable set of synthetic landscapes, we first initialize **h** and **H** by training Potts models on alignments of protein sequences. We choose protein targets to span a range of functions, structural folds, and primary sequence lengths, while ensuring the resulting Potts model has excellent contact accuracy on a high resolution structure. From the 748 Potts models trained in [63], the models corresponding to the 5 PDB IDs in Figure 3 were selected manually from the top performing models with respect to contact prediction accuracy. See Supplementary Figure S1 for predicted contact maps. We set the wildtype sequence **x**_0_ to be the alignment query sequence.

**Fig. 3:**
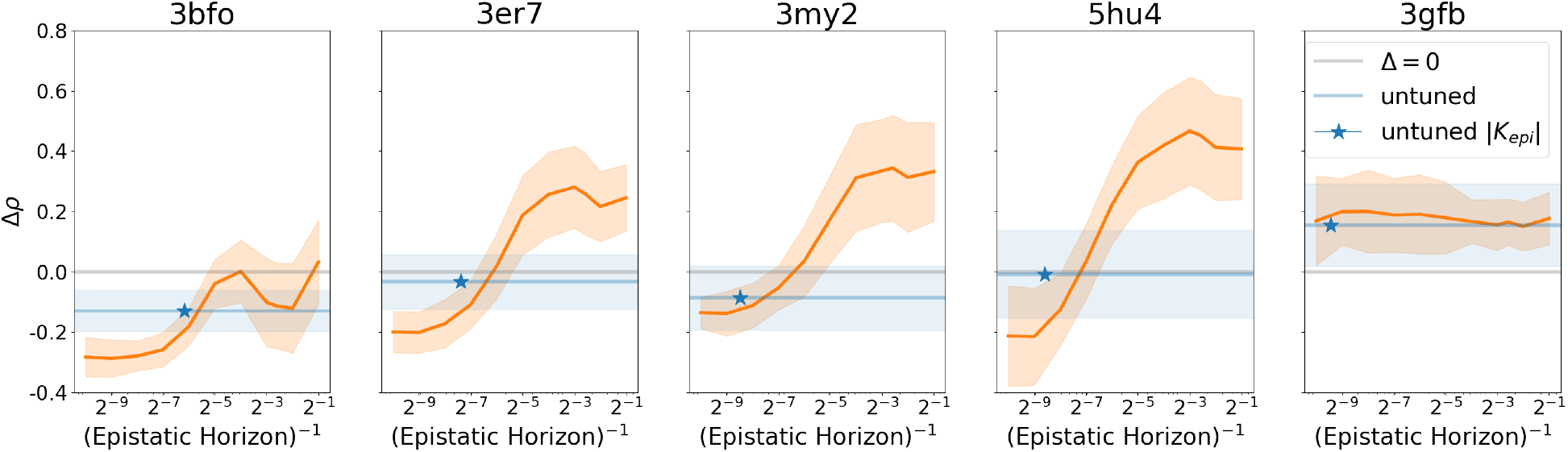
Tuning the epistatic horizon interpolates between linear and non-linear model performance on ranking combinations of 6 adaptive singles after training on 5000 examples. The grey line shows *Δρ* = 0 for ease of visualization of the threshold for one model outperforming the other. The orange line shows the difference between maximum CNN Spearman’s *ρ* and maximum Ridge Spearman’s *ρ*, with error bands showing ±1 standard deviation across 20 random training set replicates. The x-axis corresponds to inverse epistatic horizon, so that more linear landscapes are to the left and more non-linear landscapes are to the right. The blue band shows model performance on the untuned landscape. The blue star shows the position on the x-axis which corresponds to the magnitude of the untuned epistatic horizon |*K*_epi_|.

### 4.2 Evaluation Sets

We choose an evaluation set relevant for the objective of sequence design. Combining top single mutations is a common strategy for proposing variants [5], and properly ranking these proposals is directly relevant to the objective of MLDE. For each (untuned) landscape, we construct evaluation sets at mutation distance 6 from the wildtype by taking the top 20 single mutants, combining them to construct variants at the desired distance, and then taking a random subset of the desired size (200). Because the set of top 20 single mutants does not change in response to tuning (i.e., single-mutant rankings are preserved by tuning), the evaluation set for a given PDB ID is fixed to the same set of 200 sequences (note that their fitness 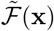 changes with tuning). See Supplementary Material A.3 for a discussion of other evaluation sets.

### 4.3 Models

#### Ridge regression

Our baseline linear model uses the `sklearn` implementation of Ridge regression, which has a single hyperparameter representing the *L*_2_ penalty. The grid of hyperparameters used during model selection is reported in Supplementary Table S3. This is a strong baseline especially for landscapes where the level of epistasis is low. In addition, Hsu *et al*. [25] showed that the ridge penalty induces a powerful inductive bias that generalizes to unseen mutations by setting the effect of unseen mutations to the average effect seen at the same position. We remove the intercept term, since centering ensures 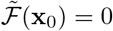.

#### Convolutional Neural Network

Convolutional neural networks (CNNs) have been used to great success in protein sequence modeling [17, 64, 65], so we select them as our nonlinear model class. The CNN model architecture is 3 layers of 1D convolutions, followed by a dense layer. On 3er7 (*L* = 118), a CNN architecture with 32 filters, kernel size 5, and hidden size 64 results in a model with 255,329 parameters. The CNN is trained using an Adam optimizer [66] to minimize MSE loss. We tune the learning rate, number of filters, batch size, and number of training epochs. See Supplementary Table S4 for tuned hyperparameters and Supplementary Table S5 for fixed hyperparameters.

### 4.4 Paired Performance Metrics

An important driver of variability in evaluation performance are the training and evaluation sets. By keeping these fixed while allowing the model class to vary, we can isolate performance differences due to modeling capacity. For each sampled training set, we select the best Ridge model and the best CNN model in terms of ranking the given evaluation set. We then compare the computed performance metric on the given evaluation set and take the difference. “Differential Spearman *ρ*” (or *Δρ*) refers to the difference, given a fixed training set, between the maximum CNN Spearman *ρ* and the maximum Ridge Spearman *ρ* on the evaluation set.

## 5 Results

### 5.1 Untuned Landscapes are Linear

In Figure 1, we plot the variant fitness as a function of the sum of constituent single-mutant effects. The untuned landscape fitness (in blue) follows a linear trend, where the fitness of a double-mutant can be well predicted by the sum of constituent single-mutant effects. Compared to the tuned landscape, the untuned landscape exhibits much less ruggedness. In Figure 3, the blue line corresponds to the differential performance of the CNN model compare to the Ridge model on the untuned landscape. Across all untuned landscapes except 3gfb, the CNN models have a mean performance change *Δρ <* 0, i.e., that the CNN does not significantly boost performance on ranking combinations of adaptive singles compared to the Ridge model. This shows that untuned Potts models do not provide landscapes useful for benchmarking sequence design guided by nonlinear models, motivating the development of our tuning framework.

### 5.2 Epistatic Horizon Tunes the Nonlinearity of the Landscape

We aim to validate that our tuning framework can create fitness landscapes difficult enough to require nonlinear models. In Figure 3, for four of the five PDB IDs, as the epistatic horizon increases (to the left in the figure), evaluation set performance skews in favor of the Ridge model, confirming that as the landscape is tuned to be more linear, linear models are preferred to nonlinear models. Conversely, as the epistatic horizon decreases (to the right in the figure) and the landscape becomes dominated by nonlinear effects, evaluation set performance shifts to favor the CNN model. The intermediate-length proteins (3er7, 3my2, 5hu4 – the middle three panels of Figure 3), show consistent behavior, indicating that the epistatic horizon is a generalizable metric of landscape difficulty.

On the intermediate-length proteins 3er7, 3my2, and 5hu4, Spearman’s *ρ* improves for the nonlinear model by between 0.2 and 0.4. For example, 3er7 shows an improvement in evaluation set performance from 0.1 to 0.3 (Supplementary Figure S2). For comparison, the authors in [12] found that zero-shot predictors on the 4-position GB1 landscape with a Spearman’s *ρ* of 0.2 are sufficient to substantially improve sequence design. Across all landscapes, all models get worse with increased tuning, indicating that decreasing the epistatic horizon increases the landscape difficulty for nonlinear as well as linear models (see Supplementary Figure S2 for unpaired model performance). For horizons *K*_epi_ ≫ *L*, the Ridge model achieves near perfect ranking accuracy *ρ* ≈1 (see Supplementary Figure S2).

On 3gfb (far right in Figure 3), differential model performance remains roughly constant around *Δρ* = 0.2 across all tested tunings. On 3bfo (the first panel of Figure 3), the shortest protein, differential model performance favors linear models for increased epistatic horizons, but does not achieve a regime where nonlinear models significantly outperform linear models. This may be due to CNN models overfitting on the short protein.

## 6 Discussion and Future Work

Using nonlinear models to guide sequence design has made a large impact in practice [13,14,17,67], but many questions still remain in regard to how to use these models as part of an MLDE pipeline. Our experimental results validate that our quadratic landscape tuning framework can generate synthetic landscapes which require nonlinear models for effective optimization. By deriving our landscapes from Potts models trained on real alignments, we ground the properties of our synthetic landscape in the structural and biochemical features of real proteins. These two properties enable tuned quadratic landscapes to be used to benchmark machine learning-guided protein design.

Our landscape tuning procedure relied on an optimization-motivated definition of the epistatic horizon *K*_epi_. There are other scenarios where a more general definition for *K*_epi_ may be more relevant; for example, in a regression setting, the relevant crossover may be the point at which the *variance* in nonlinear fitness components is more than the variance of the linear components. Additionally, we focused on a tuning which increased the difficulty of combining adaptive mutants; there are other forms of nonlinearity which make ML-aided design more useful. One such situation is finding individually non-adaptive single mutants which combine to make adaptive mutants. Our landscape tuning framework is flexible enough to allow tuning of these types of properties.

Another frontier of ML for biological sequence design is making efficient use of labeled data with protein-specific priors provided, for example, by large language models [45, 68, 69]. There is room for model development to incorporate priors that allow sequence models such as CNNs to learn more nonlinear landscapes. By providing realistic datasets for model training, tuned quadratic landscapes are a useful sandbox environment for proposing modeling advancements that can take advantage of small datasets.

Optimizing MLDE pipelines involves more than tuning a neural network architecture. In MLDE, design choices ranging from training set curation to sequence proposal distributions can have a huge impact on the overall effectiveness of the pipeline. Benchmarking these choices against tuned quadratic landscapes would allow practitioners of MLDE to understand how to optimize their pipeline before having to collect expensive experimental data. Often, a new protein design campaign will have very specific constraints, such as assay-specific noise, or limitations on experimental throughput. Our tuned quadratic landscapes lend themselves easily to multiple extensions that allow an MLDE practitioner to impose these specific constraints and see how their pipeline performs. For a new design campaign with a sequence alignment, a bespoke synthetic landscape can be derived directly to match the structural constraints of the target. In addition, MLDE design choices can be correlated with landscape difficulty by testing across a range of tuning parameters. We hope that the benchmarks enabled by SLIP will further support the development of robust and efficient methods for biological sequence design.

## Acknowledgments

This research is supported in part by an NIH grant R35-GM134922.

## Supplementary Material

### A Potts Models and Evaluation Sets

#### A.1 Landscape Details

**Table S1:**
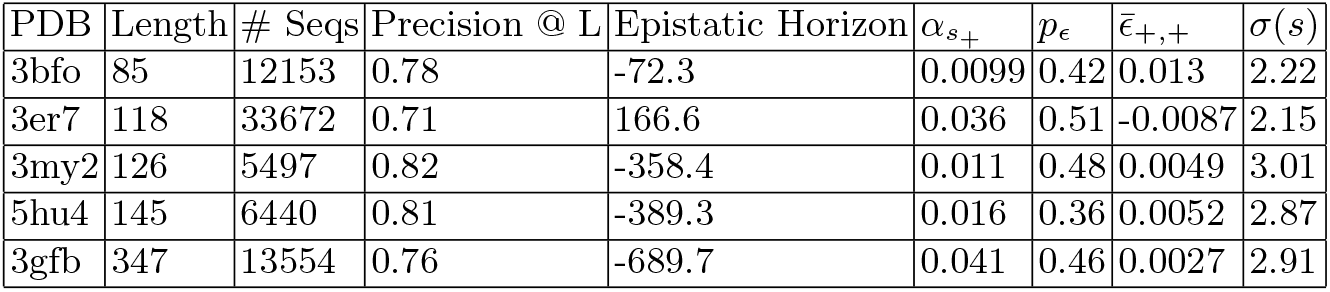
Untuned landscape details. Contact precision is computed in the standard way: predicting the top L entries (>6 apart in the primary sequence) in **H** to be contacts, and computing precision [70,71]. 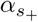 refers to the fraction of adaptive singles with effect *s*_*iβ*_ > 0. *p*_*ϵ*_ refers to the fraction of reciprocal sign epistasis for pairs of adaptive singles. 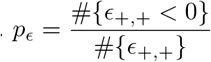

**Table S2:** Functional keywords associated with the selected PDBs.

| PDB | Functional Keywords |
| --- | --- |
| 3bfo | Immunoglobulin-like beta sandwich |
| 3er7 | Nuclear transport factor |
| 3my2 | Transmembrane protein |
| 5hu4 | Prokaryotic Sortase |
| 3gfb | L-threonine Dehydrogenase |

#### A.2 Fitting Potts Models

The initial training of the Potts model involves sampling batches from an alignment *X*. We train the model to maximize the regularized pseudolikelihood objective

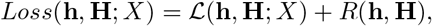

where the regularization term is given by

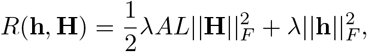

following the scaling procedure in [72]. The Potts model is trained using a modified version of Adam [66] presented in Dauparas, et al. [73], modified to improve performance of Adam to match that of L-BFGS, using batches *X*_*b*_ from the overall alignment *X*. Before computing any forward passes, we symmetrize **H** and mask the diagonal. All models were trained using *λ* = 0.01, Adam learning rate 0.5, and batch size 128. Training was done for 200 steps on a NVIDIA GeForceRTX 2080 Ti GPU. The training script can be found at https://github.com/songlab-cal/factored-attention. Models are trained using the “use-bias” flag to explicitly include **h**. From the 748 potts models trained in Bhattacharya et al. [63], the 5 PDB ids listed in Table S1 were selected manually from the top performing models with respect to contact precision @ L. Note that the deeper the alignment, the more robust the Potts model training is to hyperparameter choices.

**Table S3:** Tuned hyperparameters for the Ridge model. (Grid size: 11)

| Ridge Hyperparameter | Grid |
| --- | --- |
| L2 penalty ($\alpha$) | $10^x$ for $x \in \{-3, -2.5, -2, \dots, 1.0, 1.5, 2.0\}$ |

**Table S4:** Tuned hyperparameters for the CNN model. (Grid size: 198)

| CNN Hyperparameter | Grid |
| --- | --- |
| Learning Rate (Adam) | $10^x$ for $x \in \{-3, -2.9, -2.8, \dots, 2.0\}$ |
| Batch Size | $\{64, 128\}$ |
| Num Training Epochs | $\{100, 500, 1000\}$ |
| Num filters | $\{16, 32, 64\}$ |

**Table S5:** Fixed hyperparameters for the CNN model.

| CNN Hyperparameter | Fixed Value |
| --- | --- |
| Kernel Size | 5 |
| Hidden Size | 64 |
| Activation Function | ReLU |
| Dropout probability | 25% |

#### A.3 Epistasis-enriched evaluation sets

In Section 4.2 we describe an evaluation set based on combining multiple adaptive singles into multi-mutants. In this section we describe two additional evaluation sets for each landscape based on enrichment for epistasis. The motivation for selecting variants with large magnitude epistatic effects is to confound the linear model, and test the nonlinear model’s ability to learn epistasis from a training set of multi-mutants. We build two evaluation sets: Adaptive Epistasis and Deleterious Epistasis. For both sets the procedure for constructing them is the same, but with the sign of the epistatic terms inverted.

We construct the full *L* × *L* × *A* × *A* tensor of epistatic terms *ϵ*_*iβ,jγ*_. Ranking these terms by their value, we pick out the highest 1000 terms (for Deleterious Epistasis, we pick out the lowest 1000). From this set of strong epistatic interactions, we construct a pool of site-allele pairs: *M* = {(*iβ*_1_, *jγ*_1_), …, (*iβ*_1000_, *jγ*_1000_)}. From this pool of pairs *M* we construct variants at the desired distance from the wildtype. For an evaluation set at distance 6, we choose 3 pairs at random and combine them. We continue to sample combinations of epistatic pairs until we have an evaluation set of the desired size *n* = 200. Note that some site-allele pairs conflict with one another by mutating the same position. We discard combinations that do not reach the desired distance.

### B Quadratic Landscape Theory

In the following, we develop the basic theory for quadratic landscapes in detail. We will derive a tuning scheme which allows us to separately control the distribution of single mutant fitness effects double mutant fitness effects. This will allow us to *tune* landscapes in order to explore different regimes of the overall optimization space.

We will be interested in understanding fitness landscapes defined on sequence space. We will consider sequences of length *L* on *A* characters, encoded by vectors **x** of length *LA*, which are one-hot every *A* elements. (In computational settings, the sequence is often represented as an *L* × *A* matrix, one-hot in the second index.)

#### B.1 Quadratic fitness function

Consider a fitness function ℱ given by

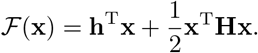

Here **H** is an *LA* × *LA* symmetric coupling matrix, and **h** is a length *LA* vector. Note that in computational settings, **H** is often implemented as a tensor of dimension *L* × *L* × *A* × *A*, and **h** as a tensor of dimension *L* × *A*.

We note that in biological applications, we generally care about fitness differences; fitness functions which differ by a constant value are considered to be the same.

Given the *L*-hot structure of **x**, we see that the *L* distinct *A* × *A*-dimensional subblocks of **H** corresponding to within-site interactions are special. In particular, only the diagonal terms contribute to ℱ, since **x** is one-hot within a block. Due to this one-hot structure, without loss of generality we can absorb the within-site interactions into the linear term by setting 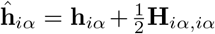 and **Ĥ** _*iα,iβ*_ = 0 for each site *i* and characters *α* and *β*, and **Ĥ** = **H** otherwise. The functional form of our fitness is unchanged:

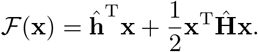

Thus, for the remainder of the notes we guarantee, without loss of generality, that in addition to being symmetric, **H** has 0 diagonal, i.e. **H**_*iα,iβ*_ = 0.

#### B.2 Local statistics

We are interested in the statistics near a particular sequence **x**_0_, which we call the “wildtype” sequence. For example, in enzyme design, we often start with a wildtype sequence which can be used in a reaction of interest, and the goal of the optimization is to arrive at a designed sequence which carries out the reaction more efficiently.

Without loss of generality, for the remainder of the notes we will refer to the wildtype sequence with a generic character **x**_0_(*i*) = *a* at all positions *i*. We often consider the relative fitness ℱ(**x**) − ℱ(**x**_0_) rather than the absolute fitness ℱ(**x**). The quadratic model is defined by two quantities: the single mutant fitness effects *s* and the *pairwise epistasis ϵ*. The single mutant fitness effects are defined by *s*(**x**_1_) = ℱ(**x**_1_) − ℱ(**x**_0_), where **x**_1_ is a single mutant which differs from **x**_0_ in exactly one of the *L* positions. We can write out the effect explicitly. Let **x**_0_(*i*) denote the character at position *i* in the wildtype sequence **x**_0_. Consider a mutation *s*_*iβ*_ at site *i*, which takes character **x**_0_(*i*) = *a* to character *β*. We have:

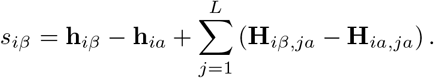

In many cases, these linear effects are enough to begin to design sequences and optimize over the fitness landscape. Note that this linear structure depends on both **h** and **H**.

The higher order interactions can be quantified using *pairwise epistasis*. In general, the term epistasis is used by geneticists to refer to interaction between the effects of multiple mutations. There are many ways to quantify these interactions. We focus on a definition of pairwise epistasis which measures the deviation from linearity of a landscape.

Given a double mutant **x**_12_, with single mutant sequences given by **x**_1_ and **x**_2_, we define the *pairwise epistasis ϵ*(**x**_12_) by

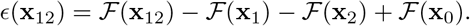

This definition can be motivated by re-writing as

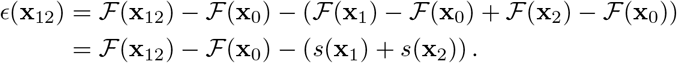

In other words, it’s the part of the fitness difference between **x**_12_ and **x**_0_ which can’t be explained by the single mutants **x**_1_ and **x**_2_.

If the two mutations are *a* → *β* at site *i* and *a* → *γ* at site *j*, then the epistasis *ϵ*_*iβ,jγ*_ is given by

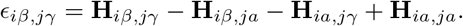

#### B.3 Difference expansion

It is useful to explicitly write the fitness difference ℱ(**x**) − ℱ(**x**_0_) for some general **x**, in order to understand and manipulate local statistics. Let *M* = {*m*_1_, *m*_2_, …*m*_*k*_} be the sequence of *k* mutations from wildtype in **x**, where *m*_*l*_ = (*i*_*l*_, *β*_*l*_). No two mutations affect the same position, so *i*_*l*_ ≠ *i*_*l*′_ for *l* ≠ *l*^′^. We have

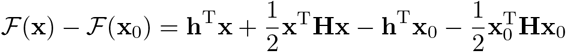

We can rewrite this in terms of the sequence difference ***δ*** ≡ **x** − **x**_0_. We have

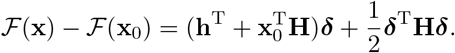

The first term captures linear effects with respect to ***δ***, and the second term captures quadratic effects. By comparing the first term to equation B.2, we can see that it in fact corresponds to the sum of the single mutant fitness effects:

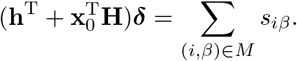

The second term is related to the epistatic effects, which we show explicitly. We have

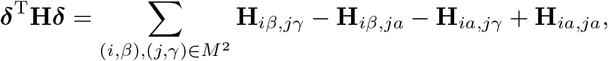

where *M*^2^ refers to ordered pairs of mutations drawn from *M*. For *i* ≠ *j*,

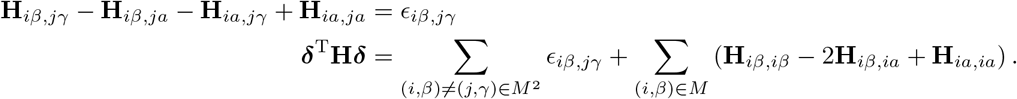

We showed that without loss of generality, we can reparameterize **H** and **h** so that **H** has no diagonal terms. Assume this is the case. Then, we can write

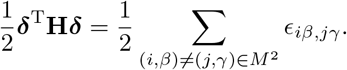

So we have shown that the second term in Equation B.3 is the sum of the pairwise epistatis effects for all pairs of mutations in **x** relative to the wildtype **x**_0_.

The expansion in Equation B.3 is useful for two reasons. Theoretically, it shows that the single mutant effects and epistatic effects control fitness differences completely, and gives us an easy way to compute them:

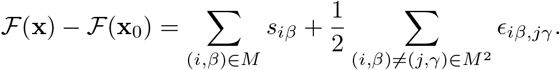

The practical consequence is that we can use the decomposition to separately manipulate the single mutant and epistasis properties of the landscape, as we will discuss in Section B.4.

#### B.4 Tuning landscapes

In order to benchmark and understand methods for exploring fitness landscapes, we want to test those methods on fitness landscapes with variable properties. In particular, given some fitness function ℱ of interest, we are interested in modifications of ℱ which make the problem “easier” or “harder” by some metric.

For a quadratic ℱ, a set of simple modifications is given by shifting and scaling the distribution of fitnesses of single and double mutants relative to the wildtype. In particular, we can independently shift (add a constant to) and scale (multipy by a constant) the single mutant statistics *s* and the epistasis statistics *ϵ* uniformly for all sequences.

As we will see, this is different from simply modifying **h** and **H**. Modifying *s* and *ϵ* corresponds to modifying the landscape in terms of first and second order expansions around the wildtype **x**_0_. In many biological problems, we care about understanding behavior near the wildtype; in addition, inferred landscape (e.g. using DCA [35]) are likely correlated with the “true” fitness landscape in limited neighborhood of the wildtype.

The shifting and scaling approach we outline maintains the relative ordering of fitnesses within the single mutants and within the double mutants. If we start with a fitness landscape whose properties are relevant for optimization, the modified landscape is one which has some similar qualitative features (e.g. important interactions in the original landscape are important in the tuned landscape). The modified landscapes can also probe different questions, such as “What happens when epistasis is more important than single mutant effects?”

We note that in most applications, we only care about the *relative* values of ℱ (e.g. ℱ(**x**) − ℱ(**x**_0_)), rather than the absolute values. We will take that approach here. If the absolute value also matters, for example if ℱ(**x**_0_) needs to be set to 0, then this can be accomplished by adding the appropriate constant to ℱ.

##### Shifting the single-mutant distribution

Suppose we wish to shift the distribution of single mutant fitness effects, relative to wildtype, by some constant *µ*_*s*_, without modifying *ϵ*. This can be accomplished by modifying **h** such that 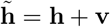. Given ℱ with parameters **h** and **H**, we define 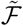 as

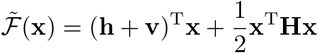

Using the expansion in Equation B.3, we have

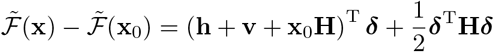

We know that the first term controls *s* and the second controls *ϵ*. Therefore, with the appropriate choice of **v**, we can modify *s* without modifying *ϵ*.

Let **v** = −*µ*_*s*_**x**_0_. We note that 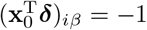 when **x** has a mutation *a* → *β* at position *i*, and 0 otherwise. Then we have:

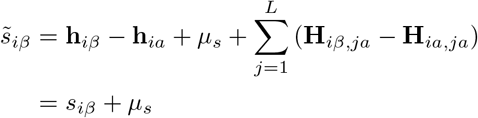

which corresponds exactly to the desired shift.

We note that the choice of **v** is not unique, since the quadratic form of ℱ, coupled with the gauge symmetry induced by the structured *L*-hot nature of of **x** means that the constant function can be written in many different ways. For example, the shift 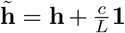 is equivalent to adding a constant *c* to the fitness function. This lack of uniqueness is not a problem computationally because our chosen form for **v** achieves the desired *s* and *ϵ* distributions - which are all that’s needed to define ℱ(**x**) − ℱ(**x**_0_).

##### Shifting the epistatic distribution

Shifting the epistatic distribution is more complicated. From Equation B.3, we see that modifying **H** affects both the epistasis distribution as well as the single-mutant distribution. Therefore, we will modify both **H** and **h** in order to modify *ϵ* without changing any of the *s*.

We shift 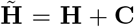 and 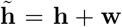. Since 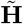 and **H** are symmetric and have 0 diagonal, **C** must be symmetric and have 0 diagonal. Our desired modified fitness function 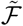 therefore, is:

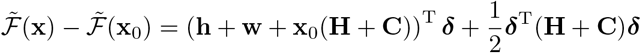

which has the same *s* as ℱ, but all *ϵ* shifted by *µ*_*ϵ*_. We proceed by deriving **C** to modify *ϵ*, and then compute **w** to ensure there is no change in *s*.

From Equation B.2, we see that one way to change the epistasis is to modify the (*ia, ja*) terms in **H**, and no others. This suggests that **C** should be proportional to 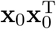, which is equal to 1 at (*ia, ja*) and 0 otherwise. We define

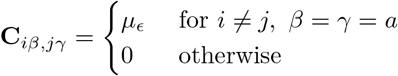

In other words,

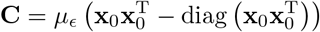

Computing the epistasis for the modified 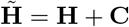 using Equation B.2, we have:

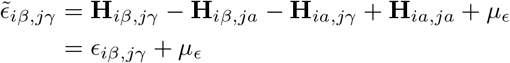

which gives us the intended shift. To ensure that *s* is unchanged, we set

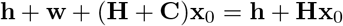

Which gives us **w** = −**Cx**_0_. Note that 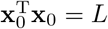 and 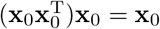. Then,

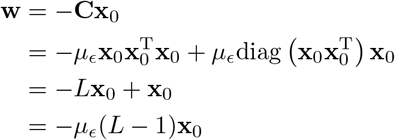

Setting **w** = −*µ*_*ϵ*_(*L* − 1)**x**_0_, *s* is left unchanged as desired.

##### Scaling the distributions

Now we consider the problem of *scaling* the distributions. That is, we want to modify **h** and **H** such that *s* and *ϵ* are multiplied uniformly by constants *λ*_*s*_ and *λ*_*ϵ*_ respectively. Using the difference expansion in equation B.3, we can see that to accomplish this we need to choose constants *A, B* and vector **u** such that:

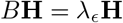

and

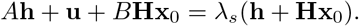

We immediately see that *B* = *λ*_*ϵ*_. In order to obtain the correct scaling of *s*, we have:

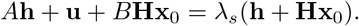

With our extra degree of freedom, we choose to set *A* = *λ*_*s*_, so we have:

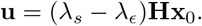

Our final fitness function is therefore

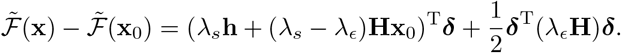

Each of the modifications outlined in Sections B.4, B.4 and B.4 can be composed:

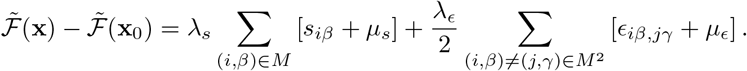

### C Supplemental Figures

**Fig. S1:**
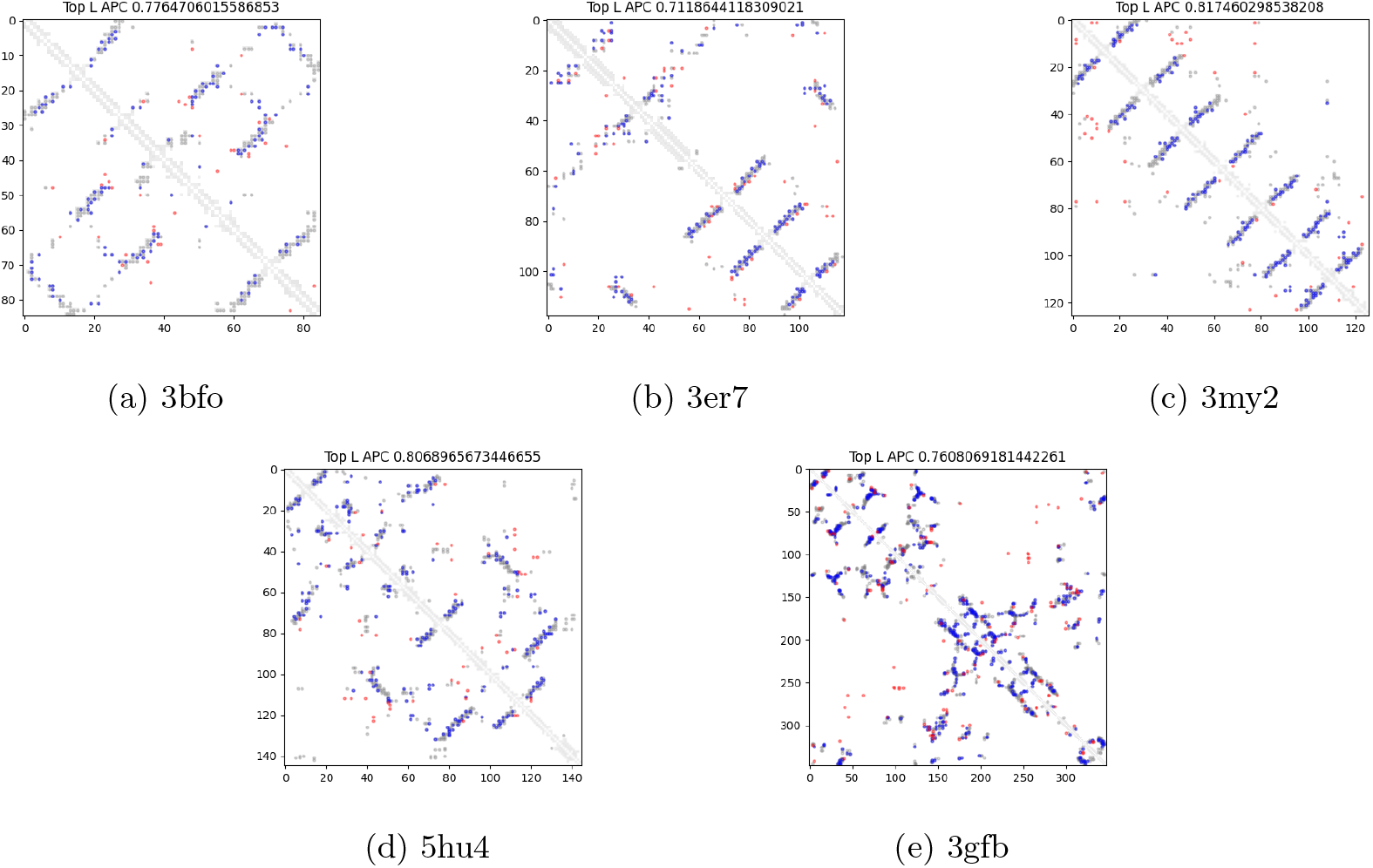
Predicted contact maps derived from Potts models fit to the corresponding alignment. Shown are Top L predicted contacts after APC correction [74], with the precision shown in the title of each plot. Blue are true predictions, red are false predictions. In grey are all contacts.

**Fig. S2:**
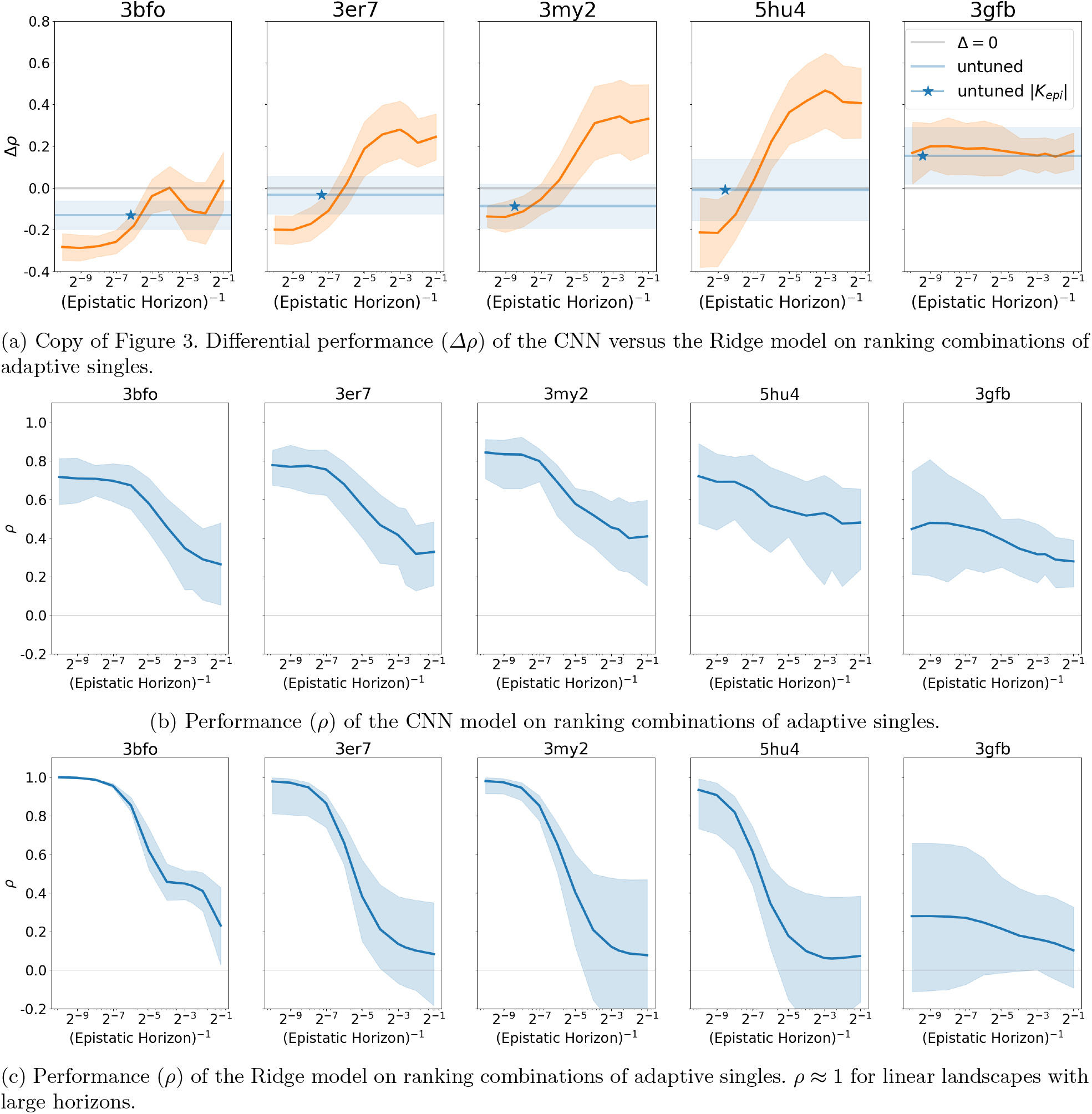
Performance (*ρ*) on ranking sequences constructed from combinations of adaptive singles.

**Fig. S3:**
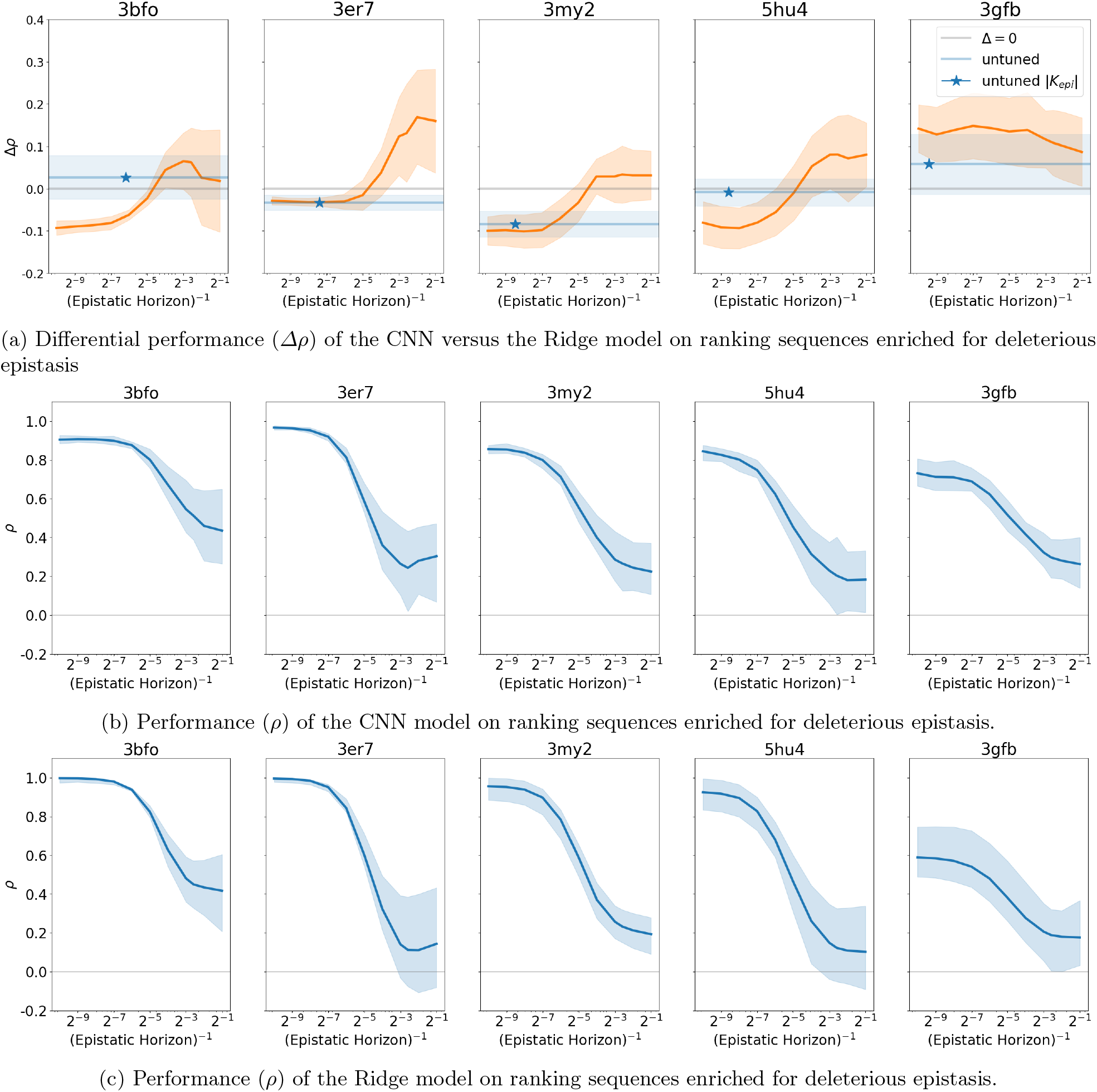
Performance (*ρ*) on ranking sequences enriched for deleterious epistasis.

**Fig. S4:**
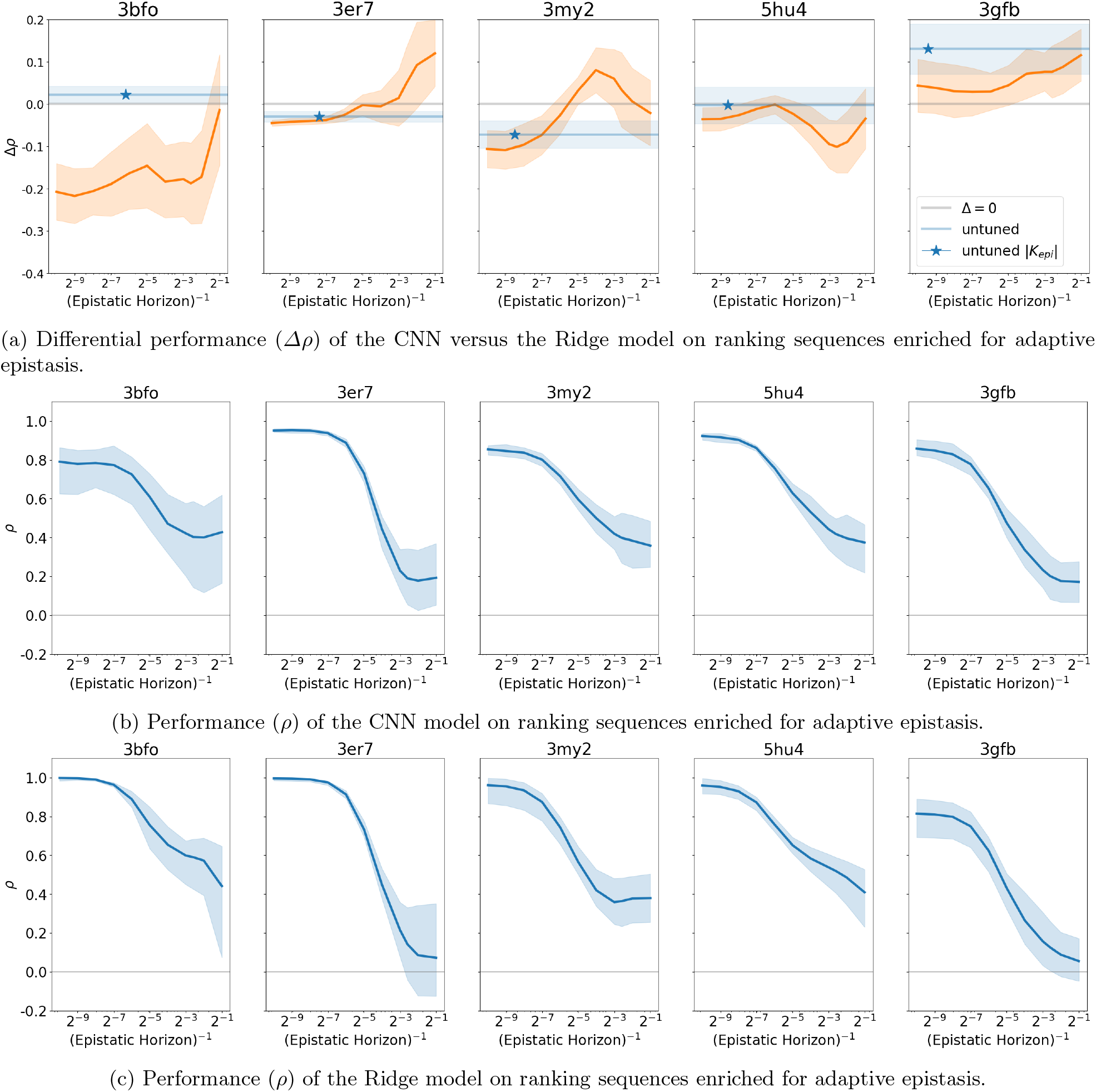
Performance (*ρ*) on ranking sequences enriched for adaptive epistasis.

**Fig. S5:**
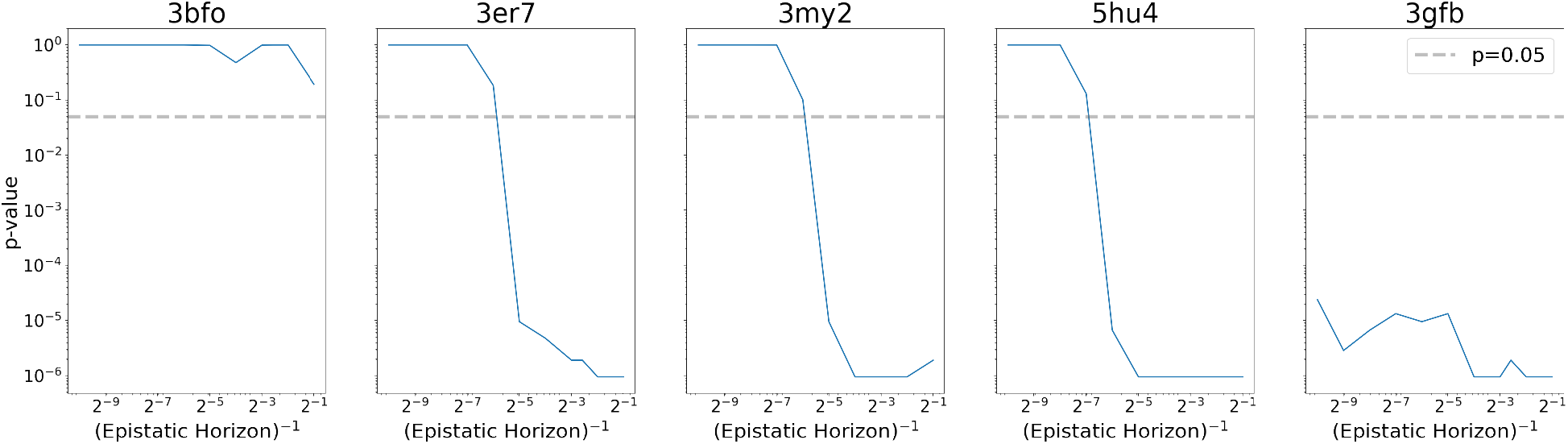
Evaluation set: Adaptive Singles. One-sided Wilcoxon signed-rank test *p*-value on *Δρ* where samples are paired by training set replicates. This tests the null hypothesis that the two models have similar evaluation performance (i.e., the distribution of *Δρ* is symmetric). The grey line indicates the significance threshold *p*=0.05. 3er7, 3my2, 5hu4 all exhibit a clear transition point where the distribution of CNN model performance is significantly better than the distribution of Ridge model performance. The CNN significantly outperforms the Ridge model across all horizons for 3gfb.

**Fig. S6:**
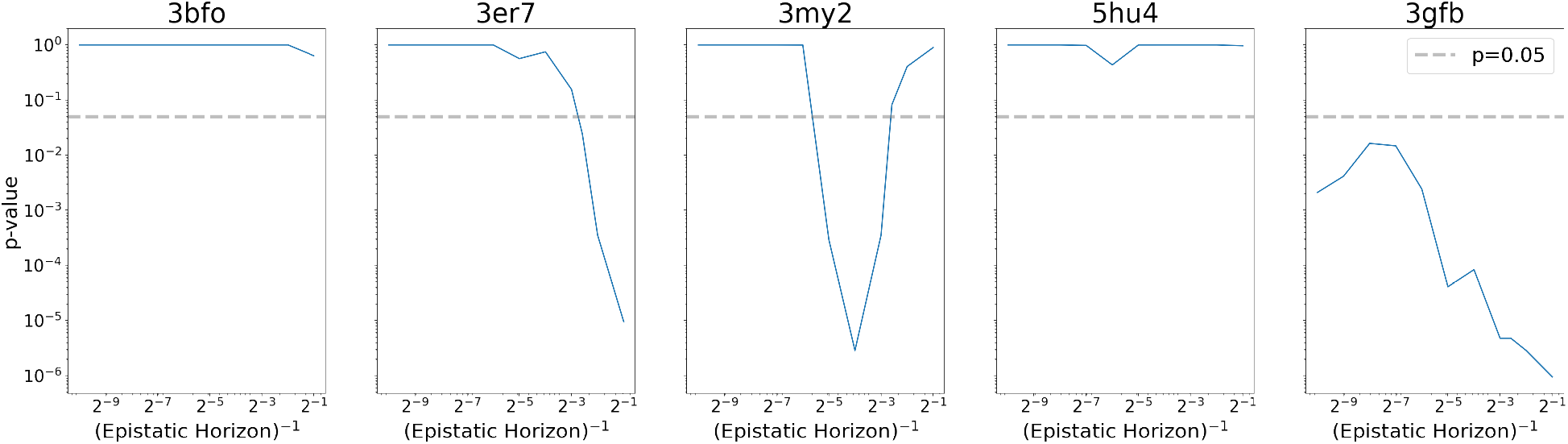
Evaluation set: Adaptive Epistasis. One-sided Wilcoxon signed-rank test *p*-value on *Δρ* where samples are paired by training set replicates. This tests the null hypothesis that the two models have similar evaluation performance (i.e., the distribution of *Δρ* is symmetric). The grey line indicates the significance threshold *p*=0.05.

**Fig. S7:**
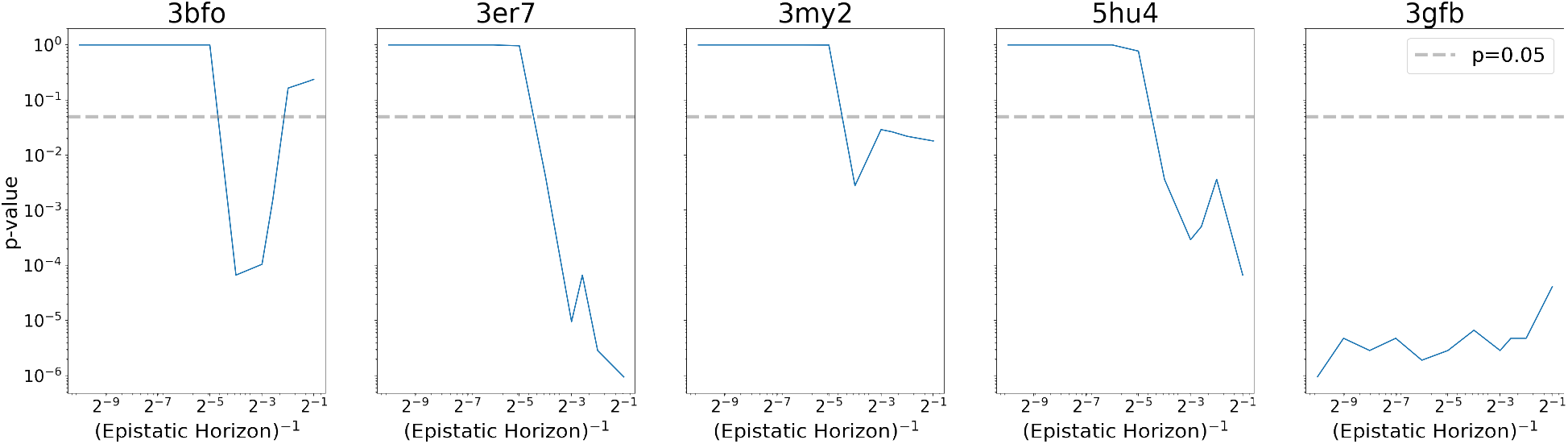
Evaluation set: Deleterious Epistasis. One-sided Wilcoxon signed-rank test *p*-value on *Δρ* where samples are paired by training set replicates. This tests the null hypothesis that the two models have similar evaluation performance (i.e., the distribution of *Δρ* is symmetric). The grey line indicates the significance threshold *p*=0.05.

**Fig. S8:**
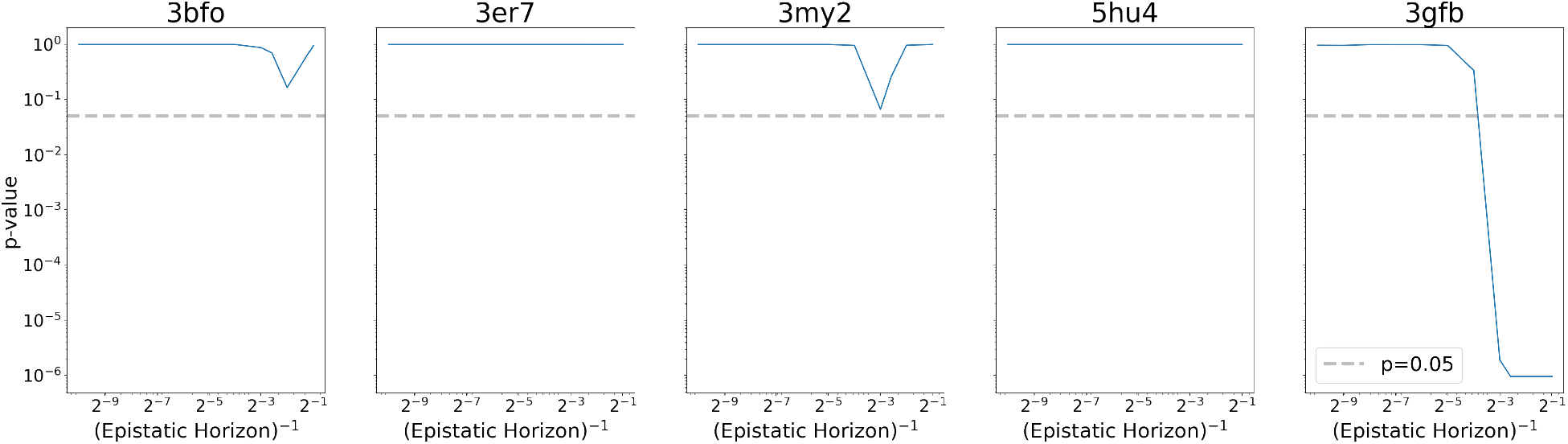
Evaluation set: Adaptive Singles. One-sided Wilcoxon signed-rank test *p*-value on *ΔMSE* where samples are paired by training set replicates. This tests the null hypothesis that the two models have similar evaluation performance (i.e., the distribution of *ΔMSE* is symmetric). The grey line indicates the significance threshold *p*=0.05.

